# ABCA1 causes an asymmetric cholesterol distribution to regulate intracellular cholesterol homeostasis

**DOI:** 10.1101/2021.12.02.471013

**Authors:** Fumihiko Ogasawara, Kazumitsu Ueda

## Abstract

Cholesterol is a major and essential component of the mammalian cell plasma membrane (PM) and the loss of cholesterol homeostasis leads to various pathologies. Cellular cholesterol uptake and synthesis are regulated by a cholesterol sensor in the endoplasmic reticulum (ER). However, it remains unclear how the PM cholesterol level is sensed. Here we show that the sensing depends on ATP-binding cassette A1 (ABCA1) and Aster-A, which cooperatively maintain the asymmetric transbilayer cholesterol distribution in the PM. ABCA1 translocates (flops) cholesterol from the inner to the outer leaflet of the PM to maintain a low inner cholesterol level. When the inner cholesterol level exceeds a threshold, Aster-A is recruited to the PM-ER contact site to transfer cholesterol to the ER. These results show unknown synergy between ABCA1 and Aster-A in intracellular cholesterol homeostasis.

## Introduction

The cholesterol transporters ATP-binding cassette A1 (ABCA1) and ABCG1 are essential for reverse cholesterol transport (RCT), a pathway through which cholesterol in peripheral tissues is delivered to the liver. ABCA1 exports cholesterol and phosphatidylcholine to apoA-I, a lipid acceptor in blood, to generate high-density lipoprotein (HDL) (Ishigami et al., 2018), and ABCG1 exports cholesterol to HDL (Kobayashi et al., 2006). ABCA1 is ubiquitously expressed in the body and HDL generation is the only pathway for RCT; thus, ABCA1 deficiency causes severe hypercholesterolemia, or Tangier disease (Bodzioch *et al,* 1999; Brooks-Wilson *et al,* 1999; Rust *et al,* 1999). Moreover, mutations in ABCA1 were found in patients with chronic myelomonocytic leukemia, suggesting that ABCA1 exerts tumor suppressor functions (Viaud et al., 2020). Regarding *in vivo* studies, the knockout of ABCA1/G1 enhances macrophage inflammatory responses (Francone *et al,* 2005; Yvan-Charvet *et al,* 2008; Zhu *et al,* 2008), and the tissue-specific knockout of ABCA1/G1 or ABCA1 has characteristic effects, including autoimmune activation in dendritic cells (Westerterp et al., 2017), impaired diet-induced obesity in adipose tissue (Cuffe et al., 2018), and less phagocytosis in astrocytes (Morizawa et al., 2017). It has been considered that these phenotypes are caused by excessive cholesterol accumulation due to defective cholesterol export. On the other hand, we recently reported that ABCA1 not only exports cholesterol, but also translocates (flops) it from the inner (IPM) to the outer leaflet of the PM (OPM) (Liu *et al*, 2017; Ogasawara *et al*, 2019; Okamoto *et al*, 2020). IPM cholesterol is maintained at around 3 mol% in various cell lines, while OPM cholesterol is 30~50 mol% (Buwaneka et al., 2021). This asymmetric cholesterol distribution allows cholesterol to function as an intramembrane signaling molecule (Liu *et al*, 2017; Ogasawara *et al*, 2020), but its physiological importance is not fully understood.

Cholesterol is synthesized in the endoplasmic reticulum (ER) and is also taken up as low-density lipoprotein (LDL) via LDL receptor. Cholesterol synthesis and uptake are regulated by sterol regulatory element binding protein (SREBP) and SREBP cleavage-activating protein (Scap) (Radhakrishnan et al., 2008). The SREBP/SCAP system is controlled by the ER cholesterol level; 5 mol% is the threshold that activates or deactivates the SREBP/SCAP system depending on the cellular cholesterol level.

In addition, cholesterol is a major component of the PM, which stores 60~90% of total cholesterol in the cell (Lange *et al,* 1989; Warnock *et al,* 1993). Recently, Sandhu et al. reported that Aster proteins (GRAMD1s) localized at the ER are recruited to the PM-ER contact site (PEcs) upon an increase in the PM cholesterol level to transfer cholesterol from the PM to the ER (Sandhu et al., 2018). Aster-B was mainly expressed in steroidogenic tissues in wild-type mice, but Aster-B knockout mice showed low steroidogenesis due to low adrenal cholesterol ester storage. Wang et al. showed that liver-specific Aster-B/C silencing ameliorated fibrotic nonalcoholic steatohepatitis by decreasing the cholesterol accumulation in hepatocytes that caused the disease (Wang et al., 2020). These studies suggest that Aster-B/C contributes to cholesterol internalization in cholesterol-abundant tissues and to cellular cholesterol accumulation. On the other hand, while it has been reported that, like Aster-B/C, Aster-A is recruited to the PEcs upon an increase in the PM cholesterol level and is ubiquitously expressed, its physiological role remains unclear. Given the above reports, we hypothesized that the low IPM cholesterol level acts as a signal for cholesterol internalization by Aster-A, which allows the SREBP/SCAP system to sense the PM cholesterol level, and that ABCA1 and Aster-A cooperatively maintain the intracellular cholesterol homeostasis.

## Results

*Gramd1b* gene, which codes Aster-B, is a direct transcriptional target of sterol-responsive liver X receptors (LXRs), and Aster-B expression is induced in response to an increased cholesterol level in the cell (Sandhu et al., 2018). We discovered, however, that Aster-A expression was induced by neither serum depletion nor LXR agonists (Figure 1), indicating that the expression level of Aster-A is not changed by the cellular cholesterol level. Considering its ubiquitous expression (Sandhu *et al,* 2018), Aster-A is expected to have a different role in systemic cells from Aster-B, which incorporates cholesterol mainly in cholesterol-abundant tissues.

**Figure 1.**
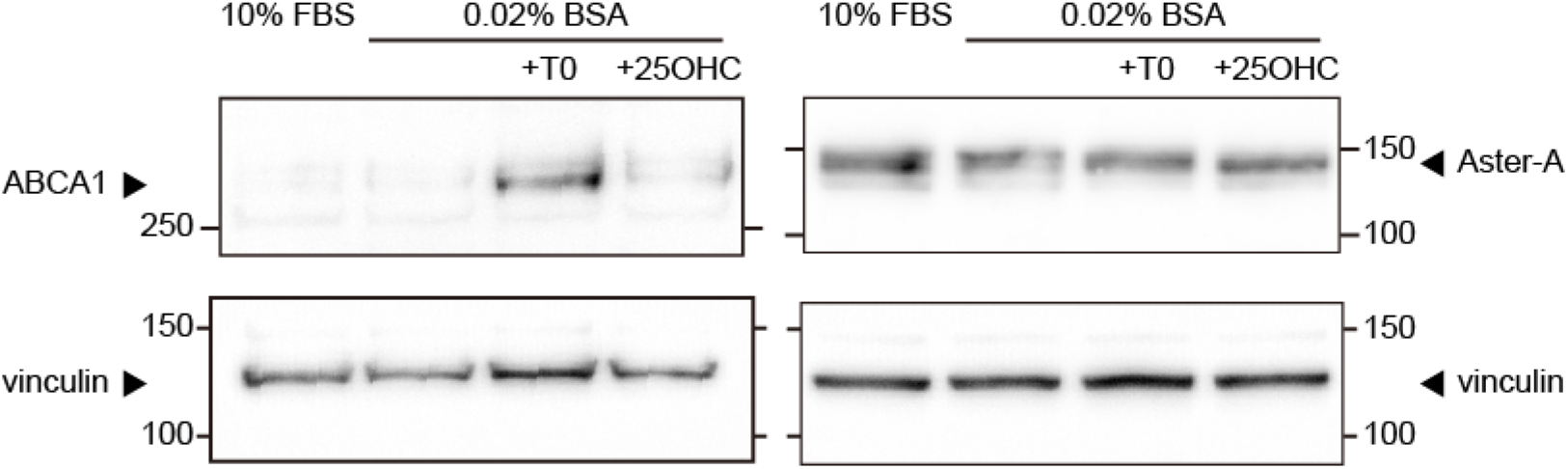
Aster-A was not a transcriptional target of LXRs. WI-38 cells, a normal human lung fibroblast line, were cultured in medium containing 10% FBS or 0.02% BSA with or without 10 μM T0901317 or 10 μM 25-hydroxycholesterol for 24 h. The expression of Aster-A, ABCA1, and vinculin, a loading control, were analyzed by western blotting.

To investigate the Aster-A function, we established HeLa cells stably expressing GFP-Aster-A. Western blotting showed a band that was shifted to a higher molecular weight than expected — the expected size is 108 kDa — but the band of endogenous Aster-A was also shifted higher (Figure 2—figure supplement 1). The band of a C-terminus deletion mutant was also shifted, but that of the Gram domain was not. Thus, the shift might be due to some modifications of the Aster domain, suggesting that GFP-Aster-A was correctly expressed. Next, the localization of Aster-A was examined by confocal microscopy. As reported (Sandhu et al., 2018), Aster-A was diffused throughout the ER and showed a peripheral dot-like distribution overlapping CellMask Deep Red, a PM marker, by cholesterol loading using the methyl-β-cyclodextrin-cholesterol complex (Figure 2a). The ratio of GFP Aster-A on the PM to that in the total cell area with and without the cholesterol loading was 0.50 and 0.16, respectively (Figure 2b). Furthermore, the movement of GFP-Aster-A on the ER network near the bottom of the cell was observed by high-resolution microscopy (Figure 2c, Movie 1). After the cholesterol loading, GFP-Aster-A molecules immediately showed a dot-like distribution in the ER network where they hardly moved for a few minutes. The dot-like distribution was localized near the PM in Figure 2a, indicating that Aster-A was recruited to the PEcs by cholesterol loading. However, when the concentration of the cholesterol loading was low, the dot-like distribution of GFP-Aster-A was dynamic, repeatedly appearing and disappearing (Movie 2). The cholesterol level of the PM does not significantly increase under physiological conditions except for some tissues or cells, such as the liver or macrophages. Thus, these results suggest that Aster-A diffuses on the ER and monitors the cholesterol level of the PM by changing the length of time it localizes at the PEcs.

**Figure 2.**
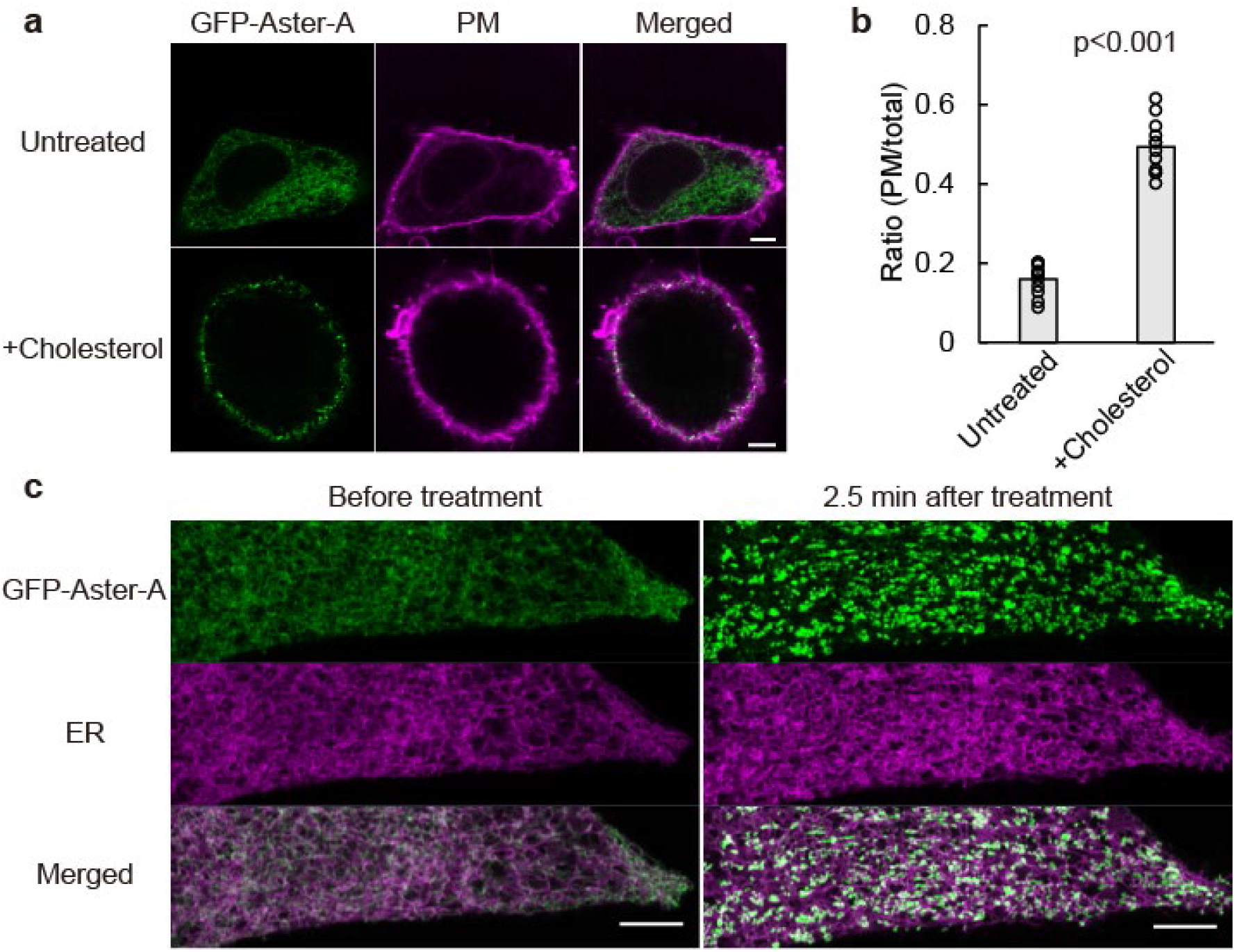
Aster-A was recruited to the PEcs by loading cholesterol. a. HeLa/GFP-Aster-A cells were treated with or without 0.4 mM MβCD-cholesterol complex for 5 min, fixed with 4% paraformaldehyde, and observed by confocal microscopy. The PM was stained with CellMask Deep Red. Scale bars, 5 μm. b. The ratio of GFP Aster-A on the PM to that in the total cell area is shown. Bars indicate average values. p < 0.001 vs. untreated. n=12 cells. c. Images of HeLa/GFP-Aster-A cells transfected with mCherry-KDEL (an ER marker) and taken using AiryScan (left). Images taken 2.5 min after a final concentration of 0.4 mM MβCD-cholesterol complex was added (right). Scale bars, 5 μm.

Previous reports show that Aster proteins sense the cholesterol concentration of the PM using their Gram domains, which bind to membranes containing cholesterol above a certain concentration (Sandhu *et al,* 2018; Naito *et al,* 2019). The Aster protein structure shows long disordered regions between the Gram domain and Aster domain, which allows the Gram domain to freely bind to IPM cholesterol. We reported that ABCA1 flops cholesterol (Liu *et al,* 2017; Ogasawara *et al,* 2019), suggesting ABCA1 suppresses Aster-A recruitment to the PEcs. To show that ABCA1 flops cholesterol, we performed flow cytometry using an Alexa Fluor 647–labeled D4 domain of perfringolysin O (PFO), a cholesterol-binding domain of pore-forming cytolysin (Figure 3a). GFP, ABCA1-GFP, or an ATP hydrolysis-deficient mutant, ABCA1(MM)-GFP, was expressed in HeLa cells. Because ABCA1 exports cholesterol to apoA-I, which is a lipid acceptor in blood and abundant in fetal bovine serum (FBS), an ABCA1 inhibitor (PSC-833) (Nagao et al., 2013) was added at time of the transfection. Before the observation, the cells were incubated in a serum-free medium for 2 hours to make ABCA1 flop cholesterol but not export it. In the cells expressing ABCA1-GFP, Alexa647-PFO-D4 binding increased with an increase in the expression level of ABCA1-GFP (Figure 3—figure supplement 1), and the median of the fluorescence intensity of Alexa647-PFO-D4 binding to ABCA1-GFP positive cells without PSC-833 increased 3.5 times compared to the condition with PSC-833 (Figure 3a). In contrast, in the cells expressing GFP or ABCA1(MM)-GFP, Alexa647-PFO-D4 binding had a negligible effect. These results showed that ABCA1 increased the OPM cholesterol level in HeLa cells. Furthermore, a decrease in the IPM cholesterol level was also examined by TIRF microscopy using PFO-D4H, which has a higher affinity for cholesterol than wild-type PFO-D4 (Maekawa and Fairn, 2015). To observe a change in the IPM cholesterol level, we established HeLa cells expressing a proper level of GFP-D4H, because the expression level of the cholesterol probe greatly affects its translocation to the PM (Buwaneka et al., 2021). When the cells were transiently transfected with ABCA1-mCherry and cultured in the presence of PSC-833, GFP-D4H gradually dissociated from the PM of the cells expressing ABCA1-mCherry after removing PSC-833 to exert ABCA1 activity, but in other cells without ABCA1 expression, GFP-D4H stayed at the PM (Figure 3b). The relative fluorescence intensity of GFP-D4H in cells expressing ABCA1 decreased to 0.26 at 4 hours, but that in cells expressing ABCA1(MM)-mCherry remained constant (Figure 3c). The same assay performed with another HeLa/GFP-D4H clone showed a similar result, suggesting that the observations are not clone-specific (Figure 3—figure supplement 2).

**Figure 3.**
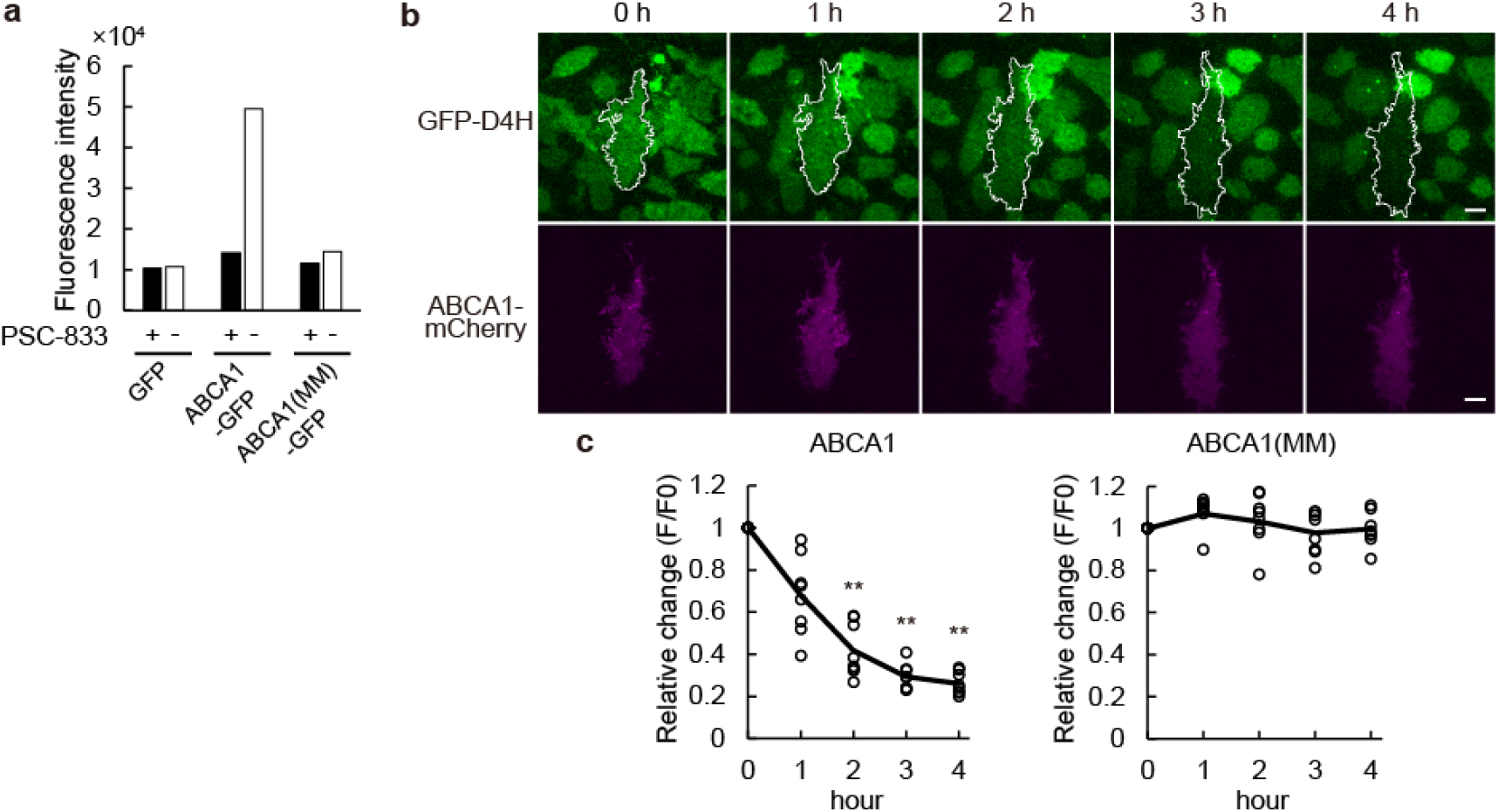
ABCA1 flops cholesterol in the PM in HeLa cells. a. HeLa cells transfected with GFP, ABCA1-GFP, or ABCA1(MM)-GFP were cultured in medium containing 10% FBS and 2.5 μM PSC-833. On the following day, the cells were incubated in serum-free medium with or without PSC-833 for 2 h, and Alexa647-PFO-D4 binding to the cells was analyzed by flow cytometry. Median fluorescence intensities of Alexa647-PFO-D4 in GFP-positive cells are shown. The flow cytometry plots are shown in Figure 3—figure supplement 1. b. HeLa/GFP-D4H cells were transfected with ABCA1-mCherry or ABCA1(MM)-mCherry in medium containing 10% FBS and 5 μM PSC-833. After the medium was changed to serum-free medium, images were acquired every hour by TIRF microscopy. The white outlines indicate the regions of the cell expressing ABCA1-mCherry. Scale bars, 10 μm. c. Relative changes in the GFP fluorescence intensity. Solid lines indicate mean values. ** p<0.001 vs. ABCA1(MM)- expressing cells. n=8-9 cells.

When cholesterol interacts with sphingomyelin in the PM, PFO cannot bind to cholesterol (Das et al., 2014). We examined whether ABCA1 increases the OPM cholesterol level even in sphingomyelin-depleted cells. Sphingomyelinase (SMase) was added at the same time PSC-833 was removed to exert the ABCA1 activity. Although SMase treatment slightly increased Alexa647-PFO-D4 binding to cells lacking ABCA1 expression, ABCA1 increased the binding regardless of SMase treatment (Figure 4a,b). SMase rapidly depleted cell-surface sphingomyelin (Figure 4c). The increase in Alexa647-PFO-D4 binding to the cells by ABCA1 confirmed the increase in the OPM cholesterol level. Therefore, these results suggest that ABCA1 flops cholesterol and decreases the IPM cholesterol level.

**Figure 4.**
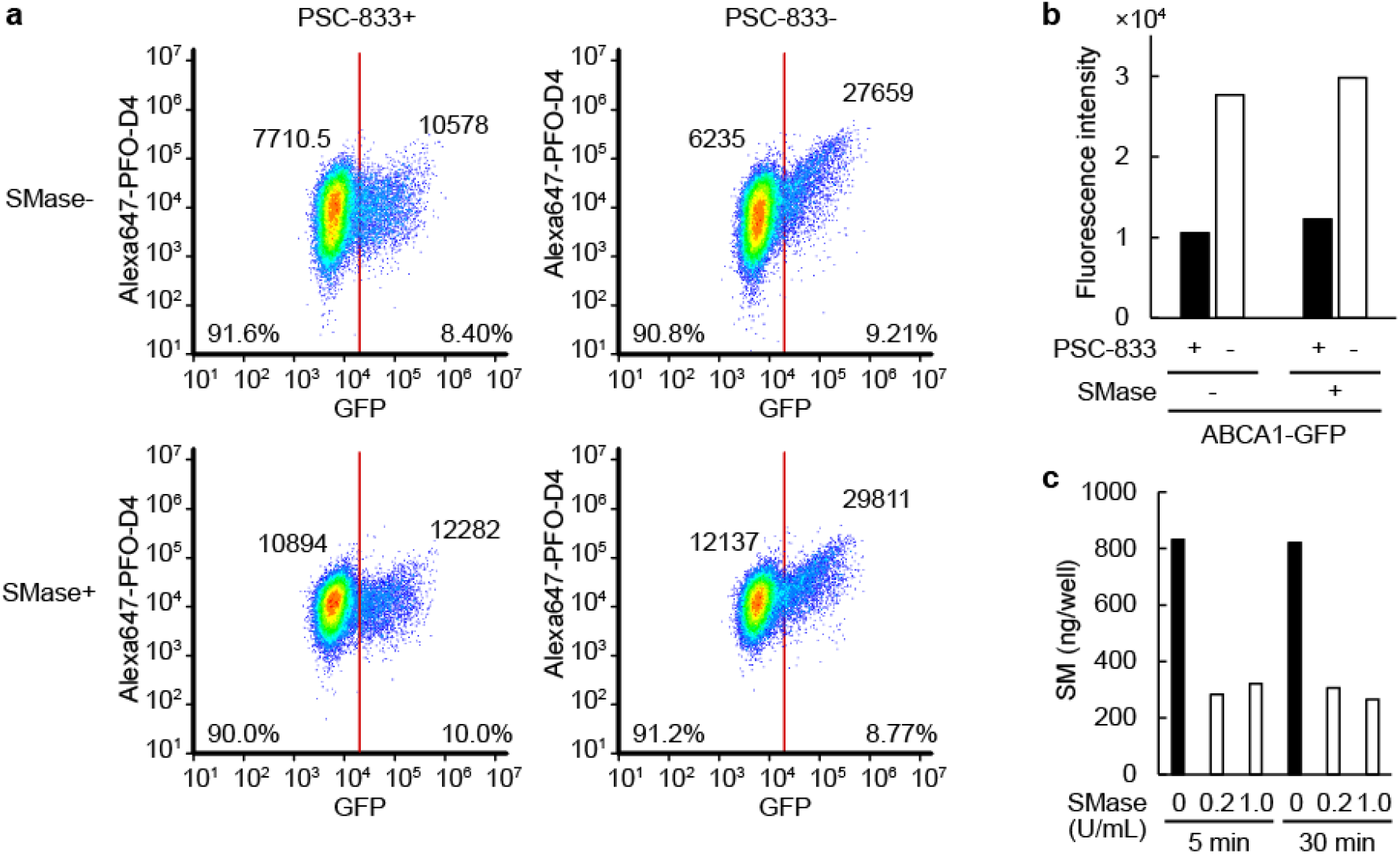
ABCA1 increased the OPM cholesterol level even in sphingomyelin-depleted cells. a,b. HeLa cells transfected with ABCA1-GFP were incubated with or without PSC-833 or SMase in serum-free medium for 2 h. Alexa647-PFO-D4 binding to the cells was analyzed by flow cytometry. (a) The plots were divided into GFP negative and positive cells at a fluorescence intensity of 20,000, and the percentage of the plots and median values of the fluorescence intensities of Alexa647-PFO-D4 are shown. (b) The median fluorescence intensities. c. The amount of sphingomyelin in HeLa/GFP-Aster-A cells was measured after SMase treatment in the indicated conditions. n=1.

Next, we examined whether ABCA1 suppresses cholesterol-dependent Aster-A recruitment. ABCA1 or ABCA1(MM) was expressed in HeLa/GFP-Aster-A cells, which were fixed at 5 minutes after cholesterol loading, and the localization of GFP-Aster-A was observed by confocal microscopy (Figure 5a). GFP-Aster-A in cells highly expressing ABCA1 was evenly diffused in the ER, but in cells expressing ABCA1(MM) it was recruited to the PEcs. The ratio of the fluorescence intensity in the PM to that in the total cell area was plotted with the ABCA1 expression level on the PM, which was measured using an antibody against the extracellular domain of ABCA1 (Okamoto et al., 2020) without permeabilization (Figure 5b). The correlation coefficient between the ratio and the ABCA1 and ABCA1(MM) expression level was −0.79 and 0.10, respectively. These plots suggested that ABCA1 suppressed cholesterol-dependent Aster-A recruitment to the PEcs depending on its expression level. To visually confirm that ABCA1 suppresses Aster-A-mediated cholesterol internalization, we added TopFluor-cholesterol, which is cholesterol conjugated with a fluorescent dye, to the PM and observed the internalization in living cells. HeLa/Aster-A cells transiently expressing ABCA1-mCherry were treated with TopFluor-cholesterol mixed with the methyl-β-cyclodextrin (MβCD)-cholesterol complex, incubated for 5 minutes, and observed by confocal microscopy (Figure 5c). However, unexpectedly, the TopFluor-cholesterol fluorescence in ABCA1-mCherry-expressing cells was lower than in cells without ABCA1 expression, and the internalization of TopFluor-cholesterol was apparently slower. Indeed, the fluorescence intensity of TopFluor-cholesterol in cells expressing ABCA1 tended to be lower in a flow cytometry assay (Figure 5d). This result was replicated in HEK293 cells, verifying the phenomenon is not cell-type specific. The high OPM cholesterol level generated by ABCA1 might prevent cholesterol transfer from the MβCD-cholesterol complex to the PM. Following these results, we decided to apply another method to examine the effect of ABCA1 on cholesterol-dependent Aster-A recruitment to test our hypothesis.

**Figure 5.**
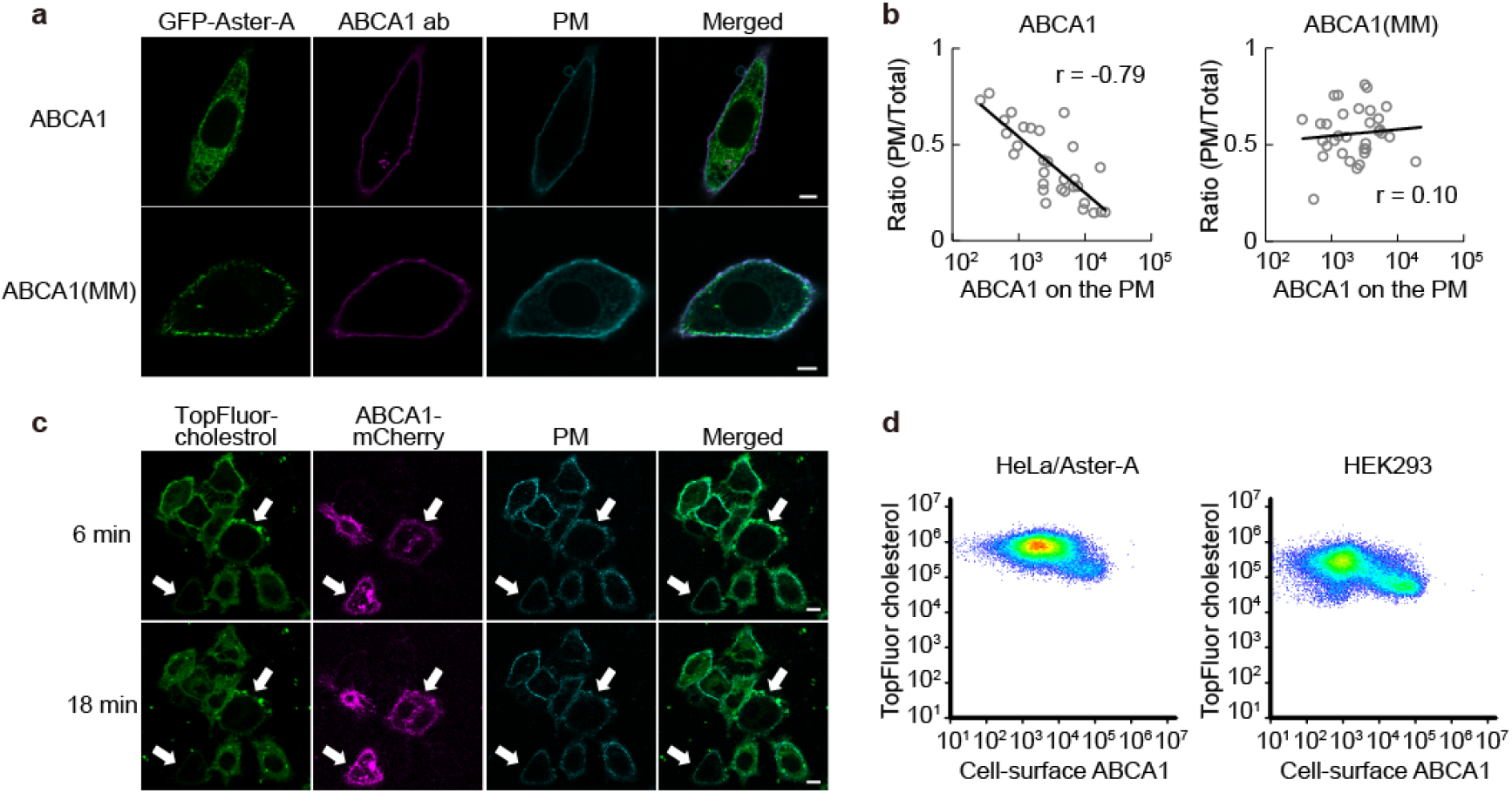
The effect of ABCA1 on cholesterol-dependent Aster-A recruitment to the PEcs. a. HeLa/GFP-Aster-A cells transfected with ABCA1 or ABCA1(MM) were treated with 0.3 mM MβCD-cholesterol complex for 5 min and fixed with 4% paraformaldehyde. ABCA1 was stained with an antibody (ab) against the anti-extracellular domain of ABCA1 and observed by confocal microscopy. Scale bars, 5 μm. b. The ratio of GFP-Aster-A on the PM to that in the total cell area is plotted with the expression level of ABCA1 or ABCA1(MM) on the PM. Log-linear regressions (solid lines) and correlation coefficients (r) are shown. n=30-33 cells. c. HeLa/Aster-A cells transfected with ABCA1-mCherry were treated with 0.2 mM MβCD-cholesterol complex mixed with Topfluor-cholesterol for 5 min and observed by confocal microscopy. White arrows indicate ABCA1-mCherry-expressing cells. Scale bars, 10 μm. d. HeLa/Aster-A cells or HEK293 cells transfected with ABCA1 were treated with 0.2 mM MβCD-cholesterol complex mixed with Topfluor-cholesterol for 5 min and analyzed by flow cytometry. Cell-surface ABCA1 was measured with ABCA1 antibody.

Sphingomyelin, which is mainly distributed at the OPM, interacts with cholesterol, but its degradation by SMase increases the IPM cholesterol level (Liu et al., 2017). Because previous reports showed that SMase treatment recruits Aster-B to the PEcs (Naito *et al,* 2019; Ercan *et al,* 2021), we used SMase to analyze the effect of ABCA1 on Aster-A recruitment. However, the effect of SMase treatment on the Aster-A recruitment was much smaller than the effect of cholesterol loading according to a confocal microscopy analysis (Figure 6—figure supplement 1). On the other hand, total internal reflection fluorescence (TIRF) microscopy clearly showed the fluorescence of GFP-Aster-A on the PEcs and a small change in the Aster-A recruitment. GFP-Aster-A formed dot-like structures near the bottom of the cells a couple of minutes after the SMase treatment (Movie 3). The mean relative fluorescence intensity reached 1.56 (Figure 6a). HeLa/GFP-Aster-A cells transiently expressing ABCA1 or ABCA1(MM)-mCherry were treated with SMase, and GFP-Aster-A was observed. GFP-Aster-A was not recruited to the PEcs in cells expressing ABCA1-mCherry (Movie 4), and the relative fluorescence intensity at ten minutes was 1.07. However, in cells expressing ABCA1(MM)-mCherry, the relative fluorescence intensity was 1.44 (Figure 6b). These results suggest that ABCA1 suppresses Aster-A recruitment to the PEcs by flopping cholesterol to the outer leaflet.

**Figure 6.**
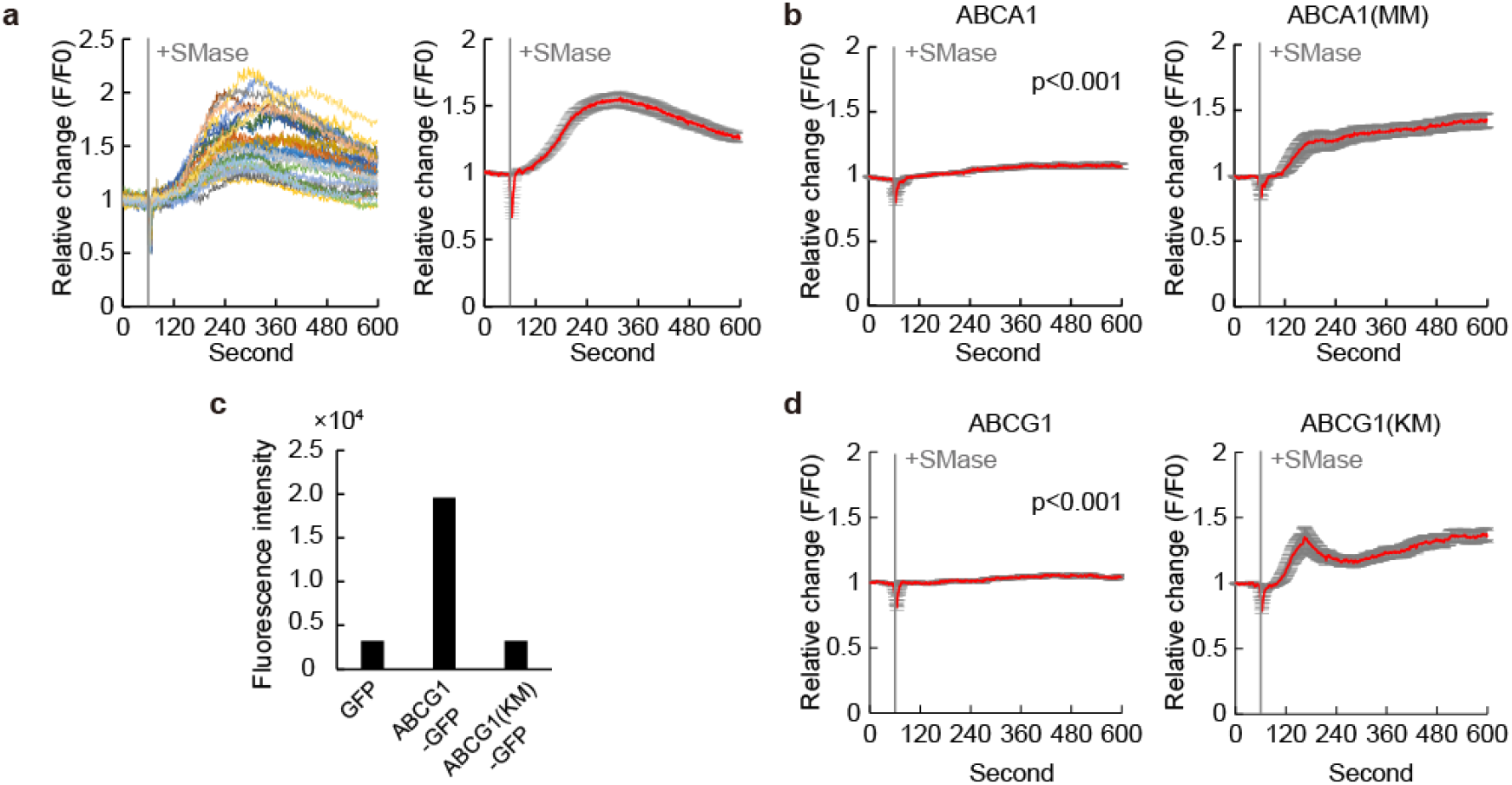
ABCA1 and ABCG1 suppressed Aster-A recruitment after SMase treatment. a. HeLa/GFP-Aster-A cells were observed by TIRF microscopy. SMase was added 60 s after the imaging began. Left, the relative change of GFP fluorescence intensity in each cell; right, mean values with S.E. n=29 cells. b. HeLa/GFP-Aster-A cells transfected with ABCA1-mCherry or ABCA1(MM)-mCherry were observed by TIRF microscopy. SMase was added 60 s after the imaging began. Mean values are shown with S.E. p<0.001 vs. ABCA1(MM)-expressing cells. The relative change in each cell is shown in Figure 6— figure supplement 2. n=24-30 cells. c. Alexa647-PFO-D4 binding to HeLa cells transfected with GFP, ABCG1-GFP, or ABCG1(KM)-GFP was analyzed by flow cytometry. The median fluorescence intensity of Alexa647-PFO-D4 in GFP-positive cells is shown. The flow cytometry plots are shown in Figure 3—figure supplement 1. d. HeLa/GFP-Aster-A cells transfected with ABCG1-mCherry or ABCG1(KM)-mCherry were observed by TIRF microscopy. SMase was added 60 s after the imaging began. Mean values are shown with S.E. p<0.001 vs. ABCG1(KM)-expressing cells at 600 s. The relative change in each cell is shown in Figure 6—figure supplement 2. n=31-33 cells.

ABCG1, which exports cholesterol to HDL (Kobayashi et al., 2006), also flops cholesterol and decreases the IPM cholesterol level (Liu *et al*, 2017; Ogasawara *et al*, 2019). To examine whether ABCG1 suppresses Aster-A recruitment to the PEcs by SMase treatment, we performed the Alexa647-PFO-D4 binding assay and TIRF microscopy. ABCG1 exports cholesterol to HDL; therefore, in the assay, FBS was removed from the medium before the transfected ABCG1 was expressed. The binding of Alexa647-PFO-D4 to cells expressing ABCG1-GFP increased with a higher expression level of ABCG1-GFP and was 6.1 times higher than in control (GFP) cells (Figure 6c). In contrast, in cells expressing ABCG1 (KM)-GFP, which is an ATP-hydrolysis deficient mutant, no significant change in Alexa647-PFO-D4 binding was observed. These findings suggest that ABCG1 flops cholesterol and decreases the IPM cholesterol level like ABCA1. The relative fluorescence intensity at ten minutes in ABCG1-mCherry expressing cells and ABCG1(KM)-mCherry-expressing cells was 1.04 and 1.34, respectively, according to the TIRF microscopy analysis (Figure 6d). These results suggest that the IPM cholesterol level is the determinant of Aster-A recruitment to the PEcs, and ABCA1 and ABCG1 maintain high OPM cholesterol and low IPM cholesterol by flopping cholesterol and suppressing Aster-A-mediated cholesterol transfer from the PM to the ER.

## Discussion

In this study, we showed that Aster-A diffuses in the ER and monitors the IPM cholesterol level by changing its length of time at the PEcs. Different methods have led to different conclusions about the asymmetric transbilayer distribution of cholesterol in the PM. However, very recently, Cho’s group showed that the IPM cholesterol level is much lower than the OPM cholesterol level by quantitative imaging analysis using two different types of ratiometric cholesterol sensors that are not appreciably affected by changes in the lipid environmental (Buwaneka et al., 2021). Some IPM cholesterol molecules are sequestered by membrane proteins such as Caveolin-1. Thus, the IPM cholesterol concentrations that make cholesterol available to cytosolic proteins, whether they are lipid sensors or signaling proteins, will also apply to Aster-A. In the ER, cholesterol is strictly maintained at 5 mol% by SREBP and Scap (Radhakrishnan et al., 2008). Therefore, IPM cholesterol above this concentration may be passively transferred by Aster-A.

In general, the SREBP-Scap system regulates cholesterol uptake, de novo synthesis, and export to maintain cholesterol homeostasis in cells (Brown & Goldstein, 1997; Horton *et al,* 2003). When the ER cholesterol level falls below the 5 mol% threshold, SREBP-2 translocates from the ER to the nucleus to induce enzymes involved in cholesterol synthesis. SREBP-2 also induces the expression of low-density lipoprotein (LDL) receptor. LDL that enters the cell via LDL receptor is degraded in lysosomes, and the bound cholesterol is delivered to the PM and then to the ER (Infante and Radhakrishnan, 2017). The knockout of Aster proteins results in cholesterol accumulation in the PM (Naito et al., 2019), and the knockdown of Aster-A specifically slows down the response of SREBP-2 to cholesterol loading to the PM (Sandhu et al., 2018). Given the above, Aster-A likely regulates cholesterol homeostasis in cells by transferring LDL-derived cholesterol to the ER. We, therefore, propose a mechanism for how the ER senses the PM cholesterol level as follows (Figure 7). First, ABCA1 flops LDL-derived cholesterol in the PM to maintain the low IPM cholesterol level. When the IPM cholesterol level exceeds a threshold, Aster-A stays at the PEcs to transfer cholesterol to the ER. The SREBP-Scap system senses the increase in the ER cholesterol level, which stops the translocation of SREBP-2 to the nucleus and thus ceases cholesterol intake and synthesis. The expression of ABCA1 also increases, which flops or exports excess cholesterol, because the suppression of ABCA1 expression by microRNA (Rayner *et al*, 2010; Najafi-Shoushtari *et al,* 2010; Horie *et al,* 2010) via SREBP-2 also ceases. The expression level of ABCA1 is immediately changed according to the ER cholesterol level because the turnover of ABCA1 is fast: 30 minutes (Azuma et al., 2009). Thus, ABCA1 and Aster-A cooperatively maintain the low IPM cholesterol level. On the other hand, Aster-A passively transfers cholesterol according to the concentration gradient (Horenkamp et al., 2018). Consequently, Aster-A has the ability to transfer cholesterol in both directions between the PM and the ER. However, Aster-A is expected to transfer cholesterol unidirectionally from the PM to the ER, since it stays at the PEcs only when the IPM cholesterol level is high. More research is needed to elucidate how cholesterol is delivered to the PM from the ER for a thorough understanding of cellular cholesterol homeostasis.

**Figure 7.**
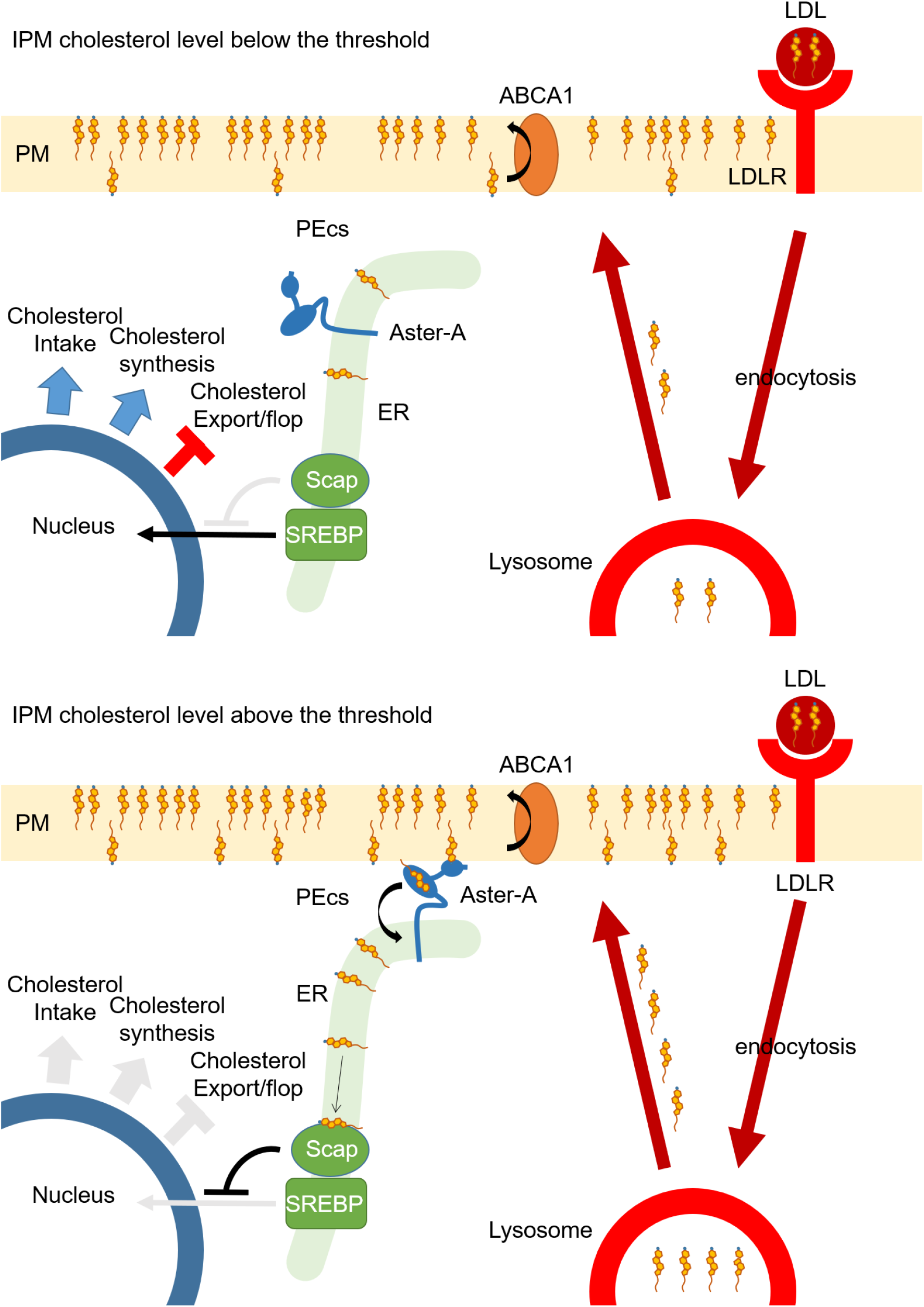
A model depicting the mechanism for how the ER senses the PM cholesterol level. Upper cartoon. ABCA1 flops LDL-derived cholesterol in the plasma membrane (PM) to maintain the low cholesterol level of the inner leaflet of the PM (IPM). Lower cartoon. As the IPM cholesterol level exceeds the threshold, Aster-A stays at the PM-ER contact site (PEcs) to transfer cholesterol to the ER. The SREBP-Scap system senses the increase in the ER cholesterol level, which stops the translocation of SREBP-2 to the nucleus and thus ceases cholesterol intake and synthesis. The expression of ABCA1 also increases, which flops or exports excess cholesterol, because the suppression of ABCA1 expression by microRNA via SREBP-2 also ceases.

Taken together, our findings show that cellular cholesterol homeostasis depends on maintaining the IPM cholesterol at 3-4 mol%, which is less than the ER cholesterol (5 mol%). This asymmetric cholesterol distribution also makes it possible for cholesterol to function as an intramembrane signaling molecule (Liu *et al*, 2017; Zhang *et al*, 2018; Ogasawara *et al*, 2020). Defective ABCA1 function prevents the asymmetric distribution of cholesterol. Consequently, signal transductions become dysregulated, resulting in the various phenotypes including cancer progression and autoimmune activation. Because the amino acid sequences of ABCA1, Aster-A, and SCAP are highly conserved among mammals, birds, and fish, the mechanism to sense the PM cholesterol level at the ER and the function of cholesterol as an intramembrane signaling molecule by the asymmetric cholesterol distribution could be common among these animals.

## Materials and Methods

### Cell culture

HeLa cells were grown in a humidified incubator (5% CO_2_) at 37°C in minimum essential medium (MEM) containing 10% heat-inactivated fetal bovine serum (FBS). WI-38 cells and HEK293 cells were grown in a humidified incubator (5% CO_2_) at 37°C in Dulbecco’s modified Eagle’s medium (DMEM) containing 10% heat-inactivated FBS.

### Plasmids

DNA insertion and site-directed mutagenesis were performed using an In-Fusion HD Cloning Kit (TaKaRa Bio) or by restriction enzyme fragmentation. The expression vectors for ABCA1-GFP, ABCA1(MM)-GFP, and ABCG1-GFP were generated as previously described (Ogasawara et al., 2019). ABCG1(KM)-GFP was generated by site-directed mutagenesis with the primers 5’-GCCGGGATGTCCACGCTGATGAACATCC-3’ and 5’-CGTGGACATCCCGGCCCCGGAAGGACCC-3’. The expression vectors for mCherry-tagged ABC proteins were generated from the expression vectors for GFP-tagged ABC proteins and mCherry cDNA by an In-Fusion reaction. The expression vector for GFP-PFO-D4 was kindly provided by Dr. Toshihide Kobayashi (University of Strasburg). PFO-D4 was generated as previously described (Ogasawara et al., 2019). D4H was generated by site-directed mutagenesis using the primers 5’-TGTTTTAGATTGATAATTTCCATCCCATGTTTT-3’ and 5’-CGGACTCAGATCTCGAAGGGAAAAATAAACTTAGA-3’ and inserted into pEGFP-C2 vector (TaKaRa Bio) by the In-Fusion reaction. GFP-D4H was then inserted into pIRESpro3 vector (TaKaRa Bio) using BamHI and NheI. Aster-A cDNA (NM_020895.5) was amplified from HEK293 cDNA by PCR using the primers 5’-TGTAAGCTTTTCGACACCACACCCCACTC-3’ and 5’-AGAGAATTCTCAGGAAAAGCTGTCATCGG-3’ and was inserted into pEGFP-C3 vector (TaKaRa Bio) using EcoRI and HindIII. GFP-Aster-A was inserted into pIRESpuro3 vector using EcoRI and NheI. ΔC mutant (1-546) and Gram domain (1-256) of Aster-A were generated by the In-Fusion reaction. mCherry-KDEL was generated by the insertion of KDEL into the mCherry C-terminus and a signal peptide into the N-terminus by the In-Fusion reaction.

### Transfection and stable cell lines

For transient expressions, cells were transfected with 1 μg/mL of each expression vector using 2 μg/mL Polyethyleneimine “MAX” (PolySciences) (Hirayama et al., 2013) in a culture medium containing 10% FBS. For stable expressions, cells were transfected with 1.25 μg/mL of each expression vector using Lipofectamine LTX (Thermo Fisher Scientific) in a culture medium containing 10% FBS and selected with 0.5 μg/mL puromycin for a couple of weeks. HeLa/GFP-D4H were cloned after the selection.

### Protein analysis

Cells were lysed with 0.1% Triton in PBS- (phosphate-buffered saline without CaCl2 or MgCl2) supplemented with EDTA-free protein inhibitor cocktail (complete, Roche) on ice. For ABCA1, proteins were diluted in sampling buffer (5 mM Tris-HCl, 40 mM DTT, 2% SDS, 1 mM EDTA, 1% sucrose, and 0.01 mg/mL pyroninY), heated at 50°C for 10 min, diluted again in sampling buffer supplemented with 5 M urea, and electrophoresed on 5%–20% PAGEL (ATTO). All other proteins were diluted in Laemmli buffer (LAEMMLI, 1970), heated at 98°C for 5 min, and electrophoresed on 5%–20% PAGEL (ATTO). The proteins were transferred to an Immobilon-P Transfer Membrane (Merck), blocked with 10% Blocking One (nacalai tesque), and blotted with the indicated primary antibody. The anti-extracellular domain of ABCA1 (MT-25) was generated as previously described (Okamoto et al., 2020). Anti-Aster-A antibody (NBP2-32148, NOVUS Biologicals), anti-vinculin antibody (V9131-2ML, SIGMA), and anti-GFP antibody (sc-9996, Santa Cruz) were purchased. Goat anti-mouse IgG (H+L) or goat anti-rabbit IgG (H+L) (Bio-Rad) were used as secondary antibodies. The immune signal was visualized using immunoStar Zeta or LD (WAKO).

### Imaging

For the confocal microscopy imaging of GFP-Aster-A, HeLa/GFP-Aster-A cells were plated in a glass-base dish (IWAKI) coated with fibronectin in MEM containing 10% FBS and incubated overnight. When transfected with ABCA1 or ABCA1(MM) (Figure 5a), the cells were plated in a 6-well plate, transfected with the indicated vectors in MEM containing 10% FBS and 2.5 μM PSC-833, and reseeded to a glass-base dish coated with fibronectin on the following day. After the cells were attached sufficiently, they were incubated in MEM supplemented with 0.02% BSA for 2 h to activate ABCA1 cholesterol-flopping activity. The cells were then treated with MβCD-cholesterol complex at the indicated concentrations or 0.2 U/mL SMase (s7651-50UN, SIGMA) for 5 min and fixed with 4% paraformaldehyde. To prepare 4 mM MβCD-cholesterol complex, 4 mM cholesterol and 36 mM MβCD were mixed in PBS- and incubated at 60°C overnight. Because ABCA1 localizes not only on the PM but also on endosomes, the expression level of ABCA1 on the PM was measured using the anti-extracellular domain of ABCA1 antibody (MT-25) with no permeabilization. The cells were blocked with 10% goat serum (SIGMA) for 30 min and stained with MT-25 and Alexa555-conjugated goat anti-Mouse IgG (H+L) (Thermo Fisher Scientific) for 1 h. The PM was stained with CellMask Deep Red (Thermo Fisher Scientific) for 10 s before observation. Imaging was performed with a LSM 700 confocal microscope equipped with α Plan-Apochromat 63x/1.40 Oil DIC M27 objective lens (Carl Zeiss).

To image the internalization of TopFluor-cholesterol, HeLa/Aster-A cells were plated in a glass-base dish coated with fibronectin, transfected with ABCA1-mCherry in MEM containing 10% FBS and 5 μM PSC-833, and incubated overnight. The cells were then incubated in FluoroBrite DMEM (Thermo Fisher Scientific) supplemented with 0.02% BSA, sodium pyruvate, and GlutaMAX (Thermo Fisher Scientific) for 1.5 h and with anti-Na+/K+ ATPase β3 subunit antibody (ECM Biosciences) and Alexa633-conjugated goat anti-Mouse IgG (H+L) (Thermo Fisher Scientific) for 30 min, and treated with 0.2 mM MβCD-cholesterol complex mixed with Topfluor-cholesterol for 5 min. Imaging was performed at 37 °C under 5% CO_2_ with the LSM 700 confocal microscope equipped with the above lens.

For high-resolution imaging, HeLa/GFP-Aster-A cells were plated in a glass-base dish coated with fibronectin, transfected with mCherry-KDEL, and incubated in MEM containing 10% FBS overnight. The cells were treated with MβCD-cholesterol complex at the indicated concentrations in FluoroBrite DMEM supplemented with 0.02% BSA, sodium pyruvate, and GlutaMAX. Imaging was performed at 37 °C under 5% CO_2_ with the LSM 880 confocal microscope equipped with α Plan-Apochromat 100x/1.46 Oil DIC M27 Elyra objective lens and an Airyscan detector (Carl Zeiss). Images were Airyscan processed automatically using Zeiss Zen2 software.

For TIRF imaging, HeLa/GFP-Aster-A cells were plated on a glass-base dish coated with fibronectin, transfected with the indicated vectors, and incubated for 6 h. The medium was then exchanged to MEM supplemented with 0.02% BSA and incubated overnight. Imaging was performed at 37 °C under 5% CO_2_ with an ECLIPSE Ti TIRF microscope equipped with an Apo TIRF 60xC Oil objective lens. Sixty seconds after beginning the movie recording, the cells were treated with 0.2 U/mL SMase in MEM without phenol red (Thermo Fisher Scientific) supplemented with 0.02% BSA and GlutaMAX. The frame rate was set to one image per second.

### Image processing and calculation

Images were processed using Fiji software to calculate the ratio of the GFP-Aster-A fluorescence intensity on the PM to that of the total cell area. A region of the PM was detected from images positive for CellMask Deep Red, and a region of the total cell was detected from the images positive forGFP-Aster-A or CellMask Deep Red. The GFP intensities on the PM and in the total cell were measured, and the autofluorescence of the HeLa parent cells was subtracted as background. When ABCA1 was expressed, the mean ABCA1 intensity on the PM was measured, and the mean autofluorescence of the HeLa parent cells on the PM was subtracted as background.

To calculate the relative change in the GFP-D4H fluorescence intensity, a region of the cell positive for ABCA1-mCherry was detected from each image. Background was removed by the subtract background command in the Fiji software, and the GFP intensity was measured.

To calculate the relative change in the GFP-Aster-A fluorescence intensity, a region of the cell was manually selected. Background was removed by the subtract background command in the Fiji software, and the GFP intensity was measured.

### Flow cytometry analysis

PFO-D4 was purified and labelled with Alexa Fluor 647 as previously described (Ogasawara et al., 2019). For the PFO-D4 binding assay, HeLa cells were plated in a 12-well plate, transfected with the indicated vectors in MEM containing 10% FBS and 5 μM PSC-833, and incubated overnight. The cells were then incubated in MEM supplemented with 0.02% BSA for 2 h. The collected cells were incubated with 0.125 μg/mL Alexa647-PFO-D4 in HBSS at 20°C for 30 min and analyzed with an Accuri C6 flow cytometer (BD). The data were exported to Excel and divided into GFP negative and positive cells at the indicated fluorescence intensities. The percentages of the plots and the median fluorescence intensities of Alexa647-PFO-D4 were calculated. Pseudocolor plots were generated using Cytospec software for each sample, and 30,000 cells were analyzed.

For the TopFluor-cholesterol assay, HeLa/Aster-A and HEK293 cells were transfected with ABCA1 in medium containing 10% FBS and 5 μM PSC-833. The cells were incubated in serum-free medium for 2 h and treated with 0.2 mM MβCD-cholesterol complex mixed with Topfluor-cholesterol for 5 min. The collected cells were incubated with the anti-extracellular domain of ABCA1 antibody (MT-25) and Alexa555-conjugated second antibody in HBSS at 20°C for 30 min and analyzed with the Accuri C6 flow cytometer. Pseudocolor plots were generated using Cytospec software.

### Measurement of sphingomyelin

HeLa/GFP-Aster-A cells were plated in a 24-well plate. On the following day, the cells were treated with SMase at the indicated concentrations for the indicated times and trypsinized. Lipids in the cells were extracted with chloroform and methanol (2:1) and dissolved in Hanks’ Balanced Salt Solution (HBSS) supplemented with 0.1% Triton X-100 and 5 mM cholic acid. 50 μL of the lipid solution was then added to a 96-well black plate and mixed with an equal volume of enzyme mixture solution (0.4 U/mL SMase, 60 U/mL CIAP, 0.4 U/mL choline oxidase, 40 mU/mL HRP, and 50 μM AmplexUltraRed (Thermo Fisher Scientific) in HBSS). After incubation at 37°C for 30 min, the fluorescence intensity of AmplexUltraRed was measured with a microplate reader (Cytation 5, BioTek).

### Statistical analysis

The statistical significance of differences between mean values was evaluated using the unpaired t-test. All experiments were performed at least twice.

## Acknowledgements

We thank Drs. Noriyuki Kioka and Yasuhisa Kimura of Kyoto University for helpful discussion. Airyscan analysis was performed at the iCeMS Analysis Center, Institute for Integrated Cell-Material Sciences (iCeMS), Kyoto University Institute for Advanced Study (KUIAS). We also thank Dr. Fumiyoshi Ishidate for technical support. This work was supported by JSPS KAKENHI Grant Numbers 18H05269 (KU), Ono Medical Research Foundation (KU), and JSPS KAKENHI Grant Numbers 20K15449 (FO).

## Author contributions

FO and KU designed experiments. FO performed experiments and analyzed data. FO and KU wrote the manuscript.

---

**Figure 2—figure supplement 1.**
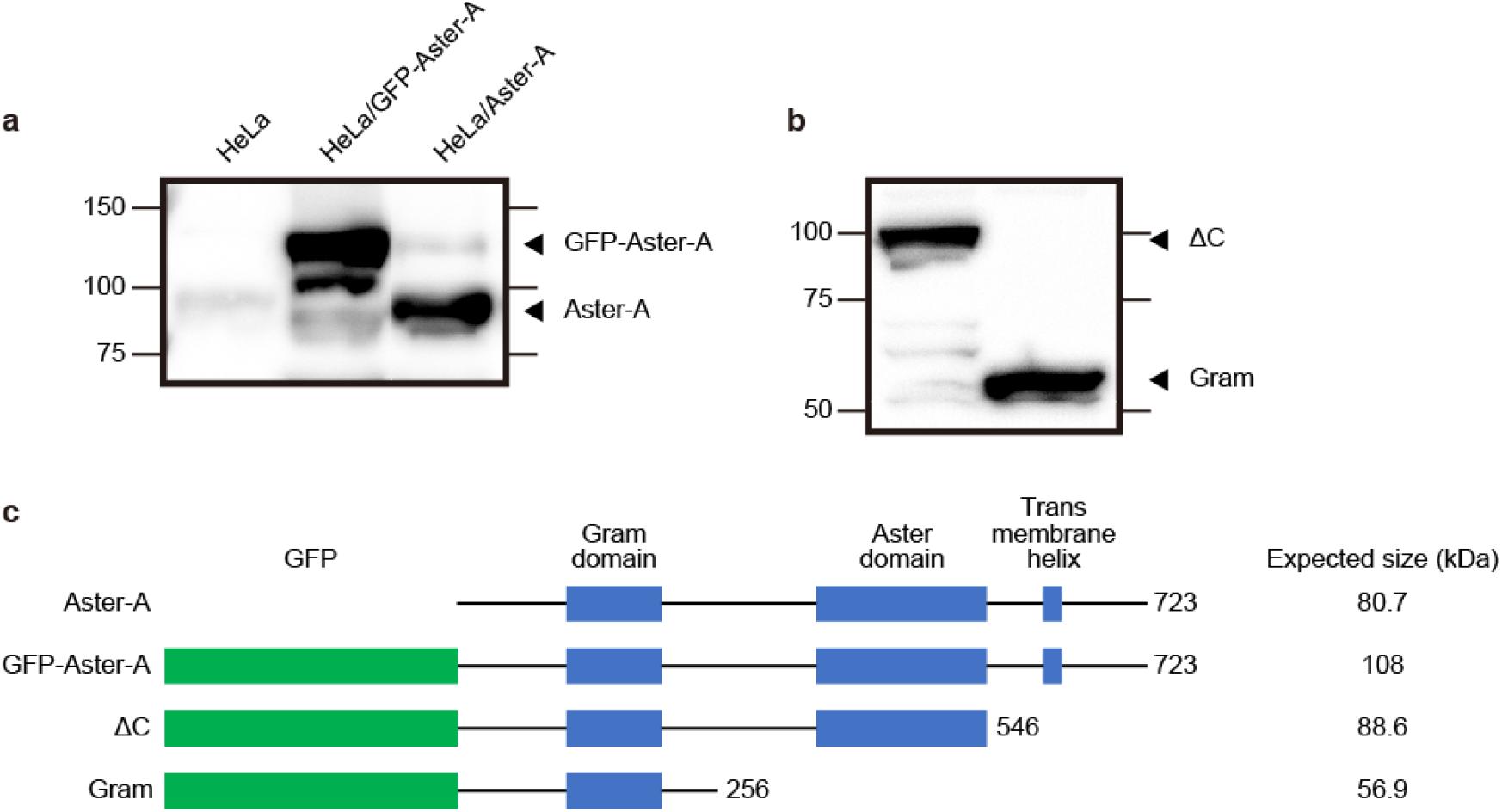
GFP-Aster-A was correctly expressed. a. The expression of Aster-A and GFP-Aster-A in HeLa cells, HeLa/GFP-Aster-A, and HeLa/Aster-A cells were analyzed by western blotting. b. ΔC mutant (1-546) and Gram domain (1-256) of Aster-A were transfected into HeLa cells, and their expressions were analyzed by western blotting with anti-GFP antibody. c. A schematic representation of Aster-A, GFP-Aster-A, GFP-Aster-A(ΔC), and GFP-Aster-A (Gram).

**Figure 3—figure supplement 1.**
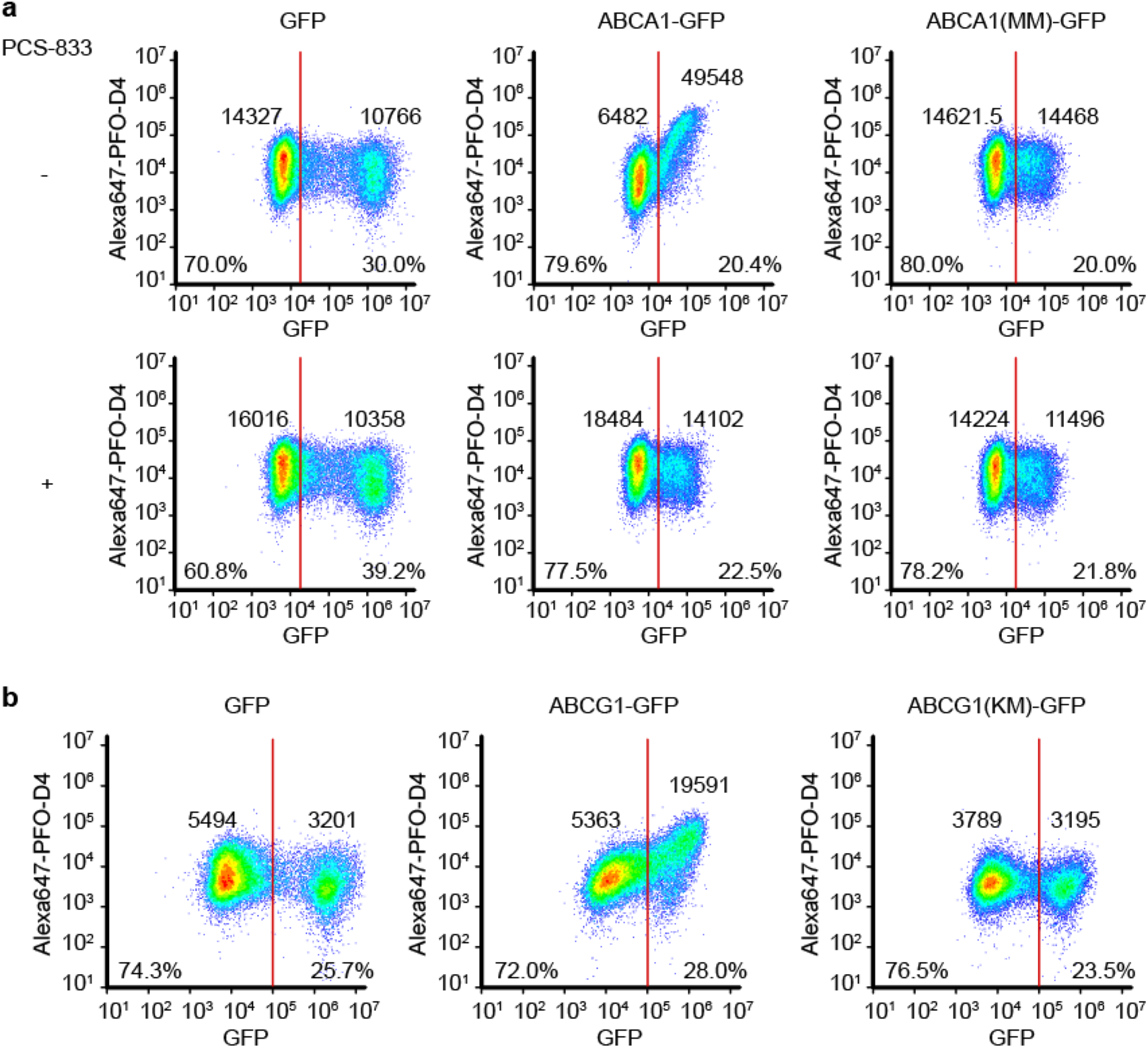
Flow cytometry plots. Flow cytometry plots of the data shown in Figure 3 (a) and Figure 6 (b). The plots were divided into GFP negative and positive cells at a fluorescence intensity of (a) 20,000 or (b) 100,000, and the percentage of the plots and median fluorescence intensities of Alexa647-PFO-D4 are shown.

**Figure 3—figure supplement 2.**
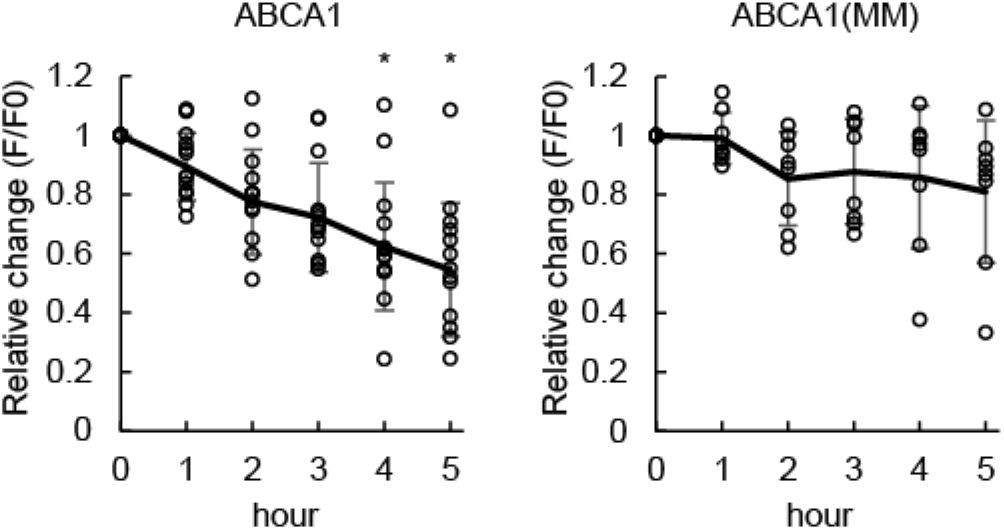
The same assay as Figure 3b,c with another clone of HeLa/GFP-D4H cells. The solid line indicates mean values. *p<0.05 vs. ABCA1(MM)-expressing cells. n=8-14 cells.

**Figure 6—figure supplement 1.**
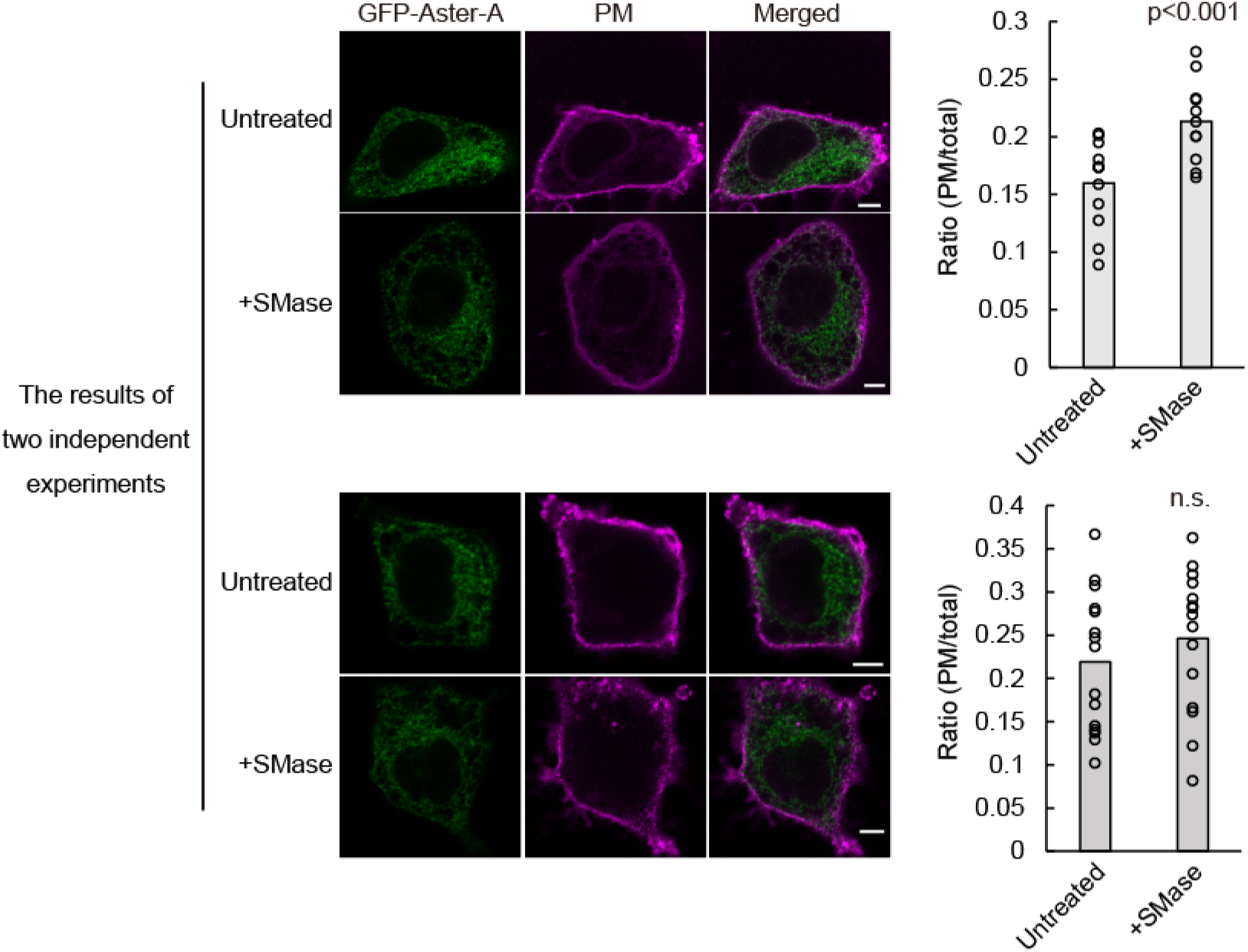
The effect of SMase on Aster-A recruitment to the PEcs was not always significant. HeLa/GFP-Aster-A cells were treated with or without SMase in serum-free medium for 5 min, fixed with 4% paraformaldehyde, and observed by confocal microscopy. The upper and lower data show the results of two independent experiments. The ratio of GFP Aster-A on the PM to that in the total cell area is shown. Bars indicate mean values. The ratio tended to increase by SMase treatment. Scale bars, 5 μm. p < 0.001 vs. untreated. n.s., not significant. n=11-15 cells.

**Figure 6—figure supplement 2.**
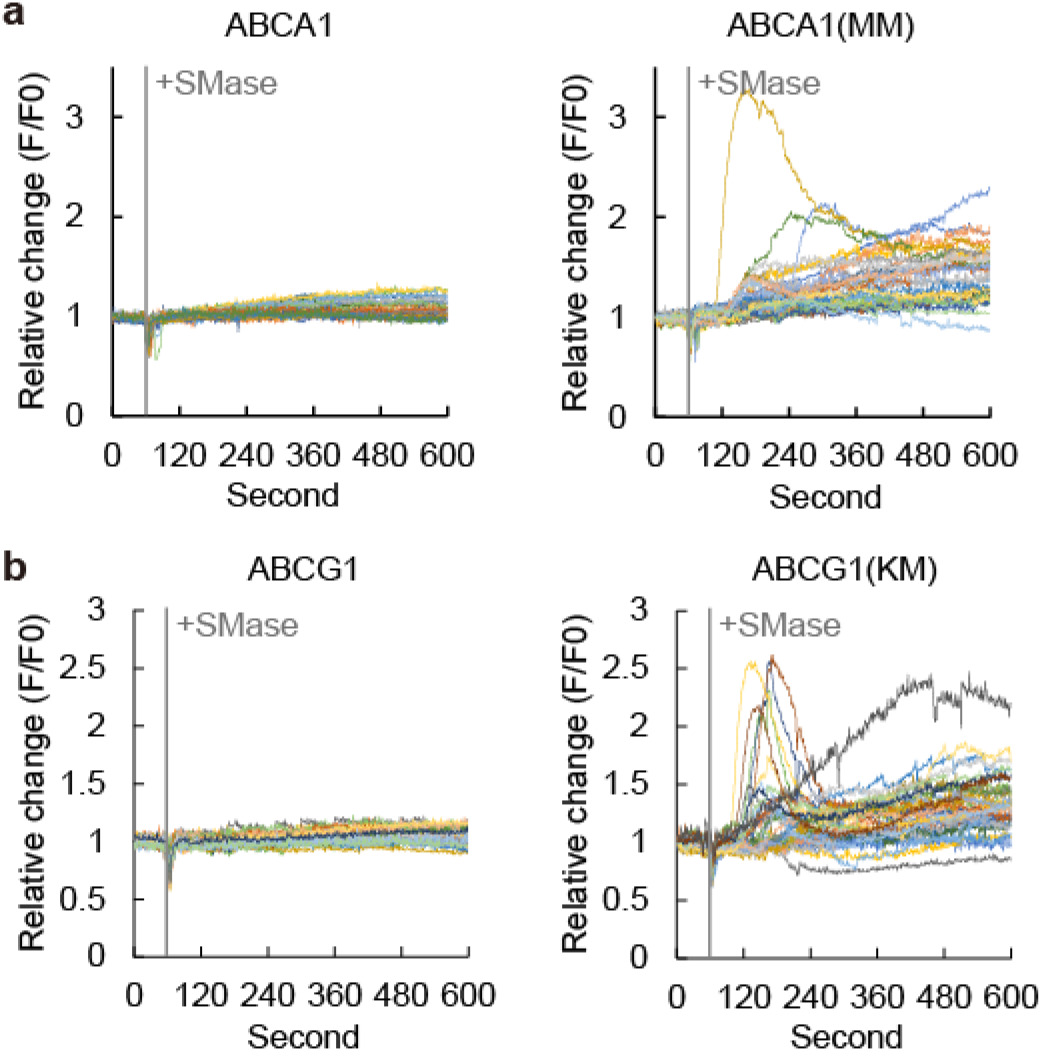
Relative change of the fluorescence intensity for each cell in Figure 6.

Movie 1. GFP-Aster-A movement after cholesterol loading.

HeLa/GFP-Aster-A cells were treated as described in the legend of Figure 2. The movie begins about 30 s after the cholesterol loading. Green, GFP-Aster-A. Magenta, ER marker (mCherry-KDEL).

Movie 2. GFP-Aster-A movement after cholesterol loading at a low concentration (0.1 mM).

The movie begins about 30 s after the cholesterol loading. Green, GFP-Aster-A. Magenta, ER marker (mCherry-KDEL).

Movie 3. GFP-Aster-A movement after SMase treatment visualized by TIRF microscopy.

HeLa/GFP-Aster-A cells were treated with SMase at 60 s.

Movie 4. GFP-Aster-A movement in cells transfected with ABCA1-mCherry after SMase treatment visualized by TIRF microscopy.

HeLa/GFP-Aster-A cells were treated with SMase at 60 s. White arrowheads show cells expressing ABCA1-mCherry.

